# Ultrasound-cell interactions mediated by cell cortex biomechanics

**DOI:** 10.64898/2026.03.25.714131

**Authors:** Dimitris Missirlis, Athanasios G. Athanassiadis, Danique Nakken, Peer Fischer

## Abstract

Low- to moderate-intensity ultrasound (US) technologies are increasingly being used to non-invasively modulate biological function in both clinical and laboratory settings. Realizing the full potential of these approaches requires a detailed mechanistic understanding of how ultrasound interacts with living cells. Here, we developed a well-controlled experimental platform to expose adherent cells to ultrasound stimulation while monitoring cellular activation via calcium imaging. We show that cell activation is dependent on cell type and identify NIH3T3 fibroblasts as a particularly robust responder. Our findings indicate that acoustic streaming is the primary mechanism underlying ultrasound-induced activation in our *in vitro* experiments. Surprisingly, the investigation of calcium dynamics revealed that the observed cytoplasmic calcium elevation originates predominantly from intracellular stores rather than extracellular influx, with membrane ion channels not contributing directly to the response. Notably, the biomechanical property of the cell-cortex emerges as a critical determinant of the cells’ sensitivity to ultrasound. Overall, our results provide clear evidence that the underlying mechanistic response involves external and internal factors that modulate the ultrasound-cell interaction and highlight important mechanistic considerations for ultrasound-based strategies aimed at cellular stimulation.

## 1. Introduction

Living cells have evolved intricate mechanisms to sense and interpret the forces they are continuously exposed to. The ability to convert compression, tensile or shear forces acting on them to biochemical information is not a feature of specialized, sensory cells, but a common cell characteristic, which naturally differs depending on cell type, tissue localization and architecture. Such information processing is possible thanks to a variety of mechanosensing and mechanotransduction mechanisms, which ultimately integrate the mechanical cues cells receive from their microenvironment to biochemical signaling pathways that cooperatively shape their phenotype and orchestrate their functional activity (Cheng et al., 2025; Saraswathibhatla et al., 2023). These mechanisms also provide an opportunity for external mechanical stimulation offers an opportunity to remotely manipulate cell activity and steer biological processes, whether in-a-dish or inside living tissue for therapeutic or intervention purposes.

Ultrasound offers an attractive approach for adjustable, non-contact, mechanical stimulation of cells, with direct relevance for clinical applications (He et al., 2025; Maresca et al., 2018; Melde et al., 2024; O’Reilly, 2024). Ultrasound comprises mechanical vibrations or pressures that oscillate in time above the range of human hearing (>20 kHz). Ultrasound waves can penetrate deep into opaque, even dense, tissues and can be focused or shaped, for instance by means of the acoustic hologram, in order to target spatially well-defined regions with high intensities, thus avoiding potentially damaging off-target effects (Jiménez-Gambín et al., 2019; Melde et al., 2016). Ultrasound does not necessarily require the introduction of additional materials to exert relevant forces, in contrast to magnetic stimulation, for instance, even though the use of ultrasound-sensitive materials may provide further control (Athanassiadis et al., 2022).

While these characteristics of ultrasound have traditionally been leveraged for ultrasound imaging or ablative therapy, a growing body of evidence indicates that cells can respond to gentler ultrasonic stimuli (He et al., 2025). Ultrasound stimulation technologies are being actively explored for clinical applications in the fields of trans-cranial neuromodulation, wound healing, and bone regeneration (Harrison et al., 2016; O’Reilly, 2024; Schandelmaier et al., 2017). Besides *in vivo* applications, substantial interest also exists to use ultrasound as a modality to guide cell proliferation or differentiation *in vitro*, including in the context of organoid research (Li et al., 2024) and micro-physiological systems (Yang et al., 2026). However, despite extensive investigations using ultrasound for remote cell stimulation, the mechanistic details remain unclear, and effective stimulation protocols remain difficult to identify. Providing a quantitative understanding on how cells decipher the exposure to ultrasound and whether the cells’ biophysical properties affect their response is much needed, as it will allow the interaction to be optimized, which in turn is needed for the efficient implementation of ultrasound stimulation technologies.

Given the millisecond timescales associated with biological responses, it is at first sight surprising that cells can even respond to MHz mechanical waves oscillating on microsecond timescales. However, it is now generally accepted that cells do not directly respond to the direct pressure oscillations of acoustic waves, but rather sense the nonlinear, or secondary, effects of the ultrasound, such as radiation forces or acoustic streaming (Kim et al., 2023; Menz et al., 2019). It is also believed that cell stimulation involves various mechano- or thermally-sensitive ion channels (Chu et al., 2022; Duque et al., 2022; Xu et al., 2023; Yang et al., 2023; Yoo et al., 2022), and it is believed that this depends strongly on the ultrasonic stimulation conditions. While a quantitative relationship between cell mechanics and acoustic stimulation has not yet been proven, it has become clear that different cell types and phenotypes respond very differently to ultrasound, suggesting an important interplay of cellular structure, mechanics, protein expression, and extracellular environment in the cellular response to ultrasound. Where precisels these secondary forces emerge and how they interact with biological structures that can differ between cell types and tissues of origin, remains, however, an open question.

Here, we set out to understand the effects of controlled ultrasound application on different cell types and identify the underlying mechanisms triggering their response *in vitro*. As a manifestation of cell activation, we opted to analyze the well-established intracellular spike of calcium ions, Ca^2+^, which act as an essential second messenger with pleiotropic effects on cells (Berridge et al., 2003). While the intracellular concentration of Ca^2+^ is maintained at low nM levels under physiological conditions, a rapid rise of cytoplasmic Ca^2+^ ions can be triggered by activating membrane ion channels present at the plasma membrane, or by release from intracellular Ca^2+^ stores at the endoplasmic reticulum. We developed an *in-situ*, focused ultrasound stimulator to monitor calcium influx in response to sonication during live-cell optical microscopy. Calcium responses were observed using small molecule fluorescent reporters, under diverse cellular microenvironments. Our experiments identified a fibroblast cell line as a strong responder, which we employed to characterize the requirements and mechanisms responsible for activation. Our results demonstrate that under these ultrasound conditions, the dominant mechanism for cell activation is acoustic streaming. Furthermore, our data reveal that the biomechanical properties of the cell cortex and plasma membrane are determining factors in the ultrasound-triggered responses.

## 2. Results

### 2.1 NIH3T3 fibroblasts respond strongly to ultrasound stimulation

We designed and built a focused ultrasound platform to apply ultrasound pulses in-situ on adherent cells cultured in a glass-bottom dish (Figure 1A). The platform consisted of a focused ultrasound transducer immobilized on a 3D-printed plastic mount, which sealed the cell culture dish and precisely positioned the ultrasound focus in the center of the dish (Figure S1). Ultrasonic stimuli consisted of repeated sinusoidal pulse trains, with a base frequency of 3 MHz, and with varying amplitudes and temporal pulse patterns. The strength and temporal profile of the ultrasonic stimulation were user-controlled via a function generator and are presented here as an effective pressure acting on the cells.

**Figure 1.**
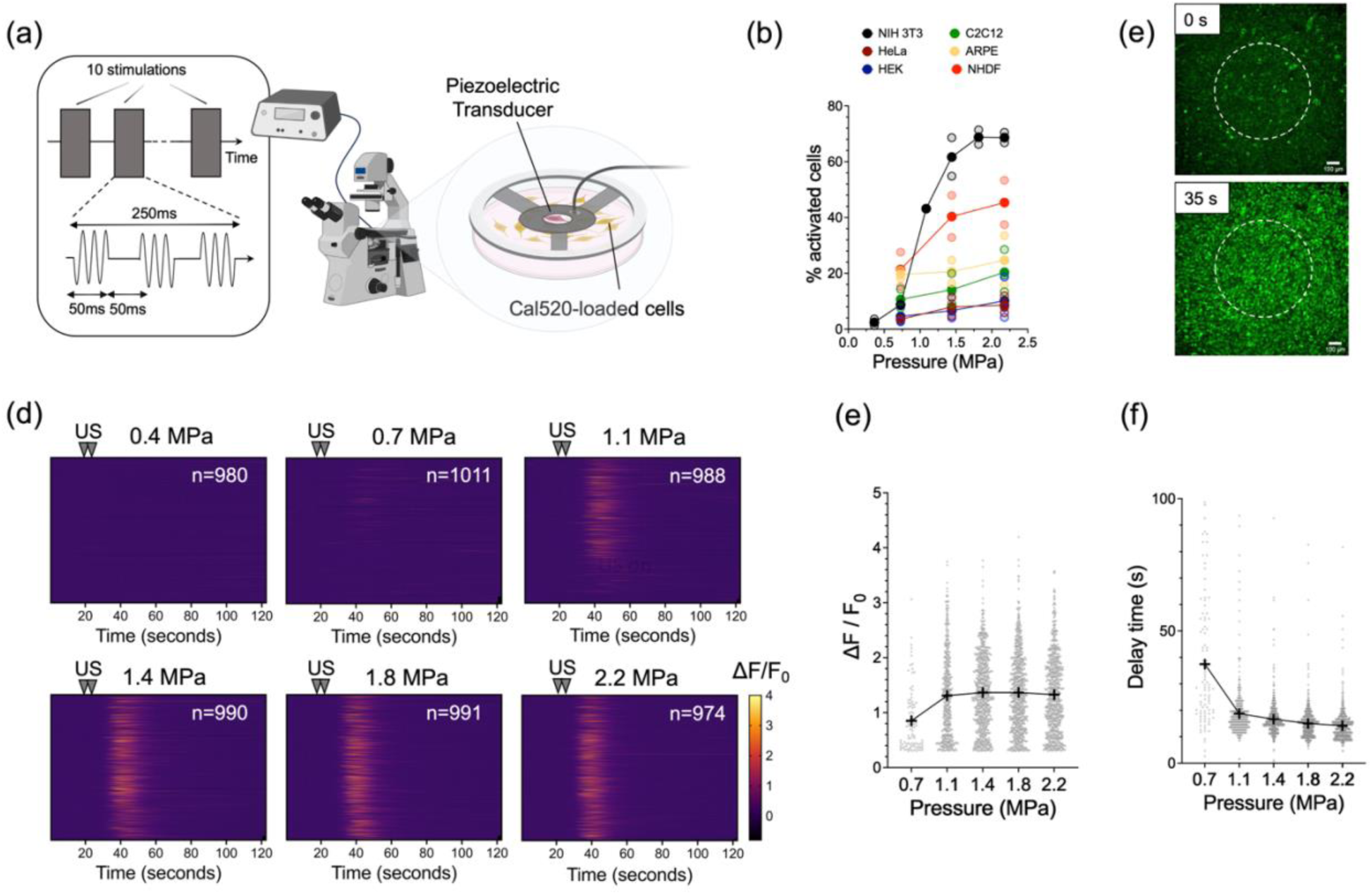
Cell response to ultrasound is cell-type dependent; NIH3T3 fibroblasts exhibit robust, pressure-dependent activation. **a)** Setup for live cell exposure to focused US with concurrent imaging. The fluorescence intensity of Cal520-loaded adherent cells in the middle of a glass-bottom dish can be recorded during exposure to an insonation profile that is controlled via a function generator. **b)** Percentage of activated cells as a function of applied pressure for the different cell types tested. Light circles correspond to independent experiments and solid circles to their average. **c)** Fluorescence microscopy images of Cal520-loaded NIH3T3 fibroblasts before and 15 seconds after sonication (35 seconds from begin of timelapse). The dashed circle denotes the US focal spot. **d)** 2D histograms for NIH3T3 fibroblast activation at the different acoustic pressures; the number of cells/ROIs is indicated for each condition. **e)** Peak normalized fluorescence intensity and **f)** time delay between peak ultrasound application and peak intensity, recorded for NIH3T3 at different acoustic pressures (each data point corresponds to an individual cell/ROI and crosses indicate the mean value).

A panel of common cell lines was screened to identify cell types that respond to focused ultrasonic waves. As a measure of cell activation, intracellular calcium levels were monitored in cells loaded with the Ca^2+^ reporter dye Cal520AM. Cells were exposed to a pulse train lasting a total of 2.5 seconds unless otherwise noted (exact pulse parameters given in Table 1 following the ITRUSST recommendations (Martin et al., 2024) and referred to as control US parameters hereafter). The control US parameters that robustly activate certain cell lines were determined in a series of preliminary experiments. The fluorescence signal was monitored for 120 seconds at 1 Hz (frame rate) that included a 20-second pre-stimulation recording to ascertain the background activity. The obtained time-lapse sequences were analyzed to determine the fluorescence intensity versus time for hundreds of individual cells (see experimental section and Figure S2 for details). Following cell segmentation, the fold-increase in dye fluorescence intensity (ΔF/F_0_) was calculated and plotted as a function of time. An activation threshold of ΔF/F_0_=0.3 was selected to calculate the fraction of activated cells and the peak intensity for each cell for the activated population (Figure S2).

**Table 1.** Ultrasound pulse timing parameters used in this study.

|  | **Duration** | **Ramp Duration** | **Ramp Shape** | **Repetition Interval/Frequency** |
| --- | --- | --- | --- | --- |
| **Pulse** | 50 ms | 0 | rectangular | 100 ms / 10 Hz |
| **Pulse Train** | 250 ms | 0 | rectangular | 500 ms / 2 Hz |
| **Pulse Train Repeat** | 2.5 s | 0 | 0 | - |

The response of cells to increasing acoustic pressures was cell type-dependent. HEK 293 cells, often used as a model to test over-expression of mechanosensitive channels, and HeLa cells, a common cancer cell line, barely responded above background levels under our experimental conditions (Figure 2b and Movie S1). The ARPE-19 retinal epithelial cell line exhibited high background level of activation, but little change upon US stimulation (Figure 2b and Movie S1). Myoblast C2C12 cells also showed spontaneous Ca2+ oscillations, which were enhanced with increasing ultrasound intensities; however, only 1/4^th^ to 1/3^rd^ of cells responded at the highest pressures examined (Figure 2b and Movie S1). In contrast, NIH3T3 fibroblasts exhibited robust and reproducible calcium influx with increasing pressure (Figure 2b and Movie S1). We also tested primary human dermal fibroblasts (NHDF), which showed a similar, albeit weaker response with increasing pressures (Figure 2b).

**Figure 2.**
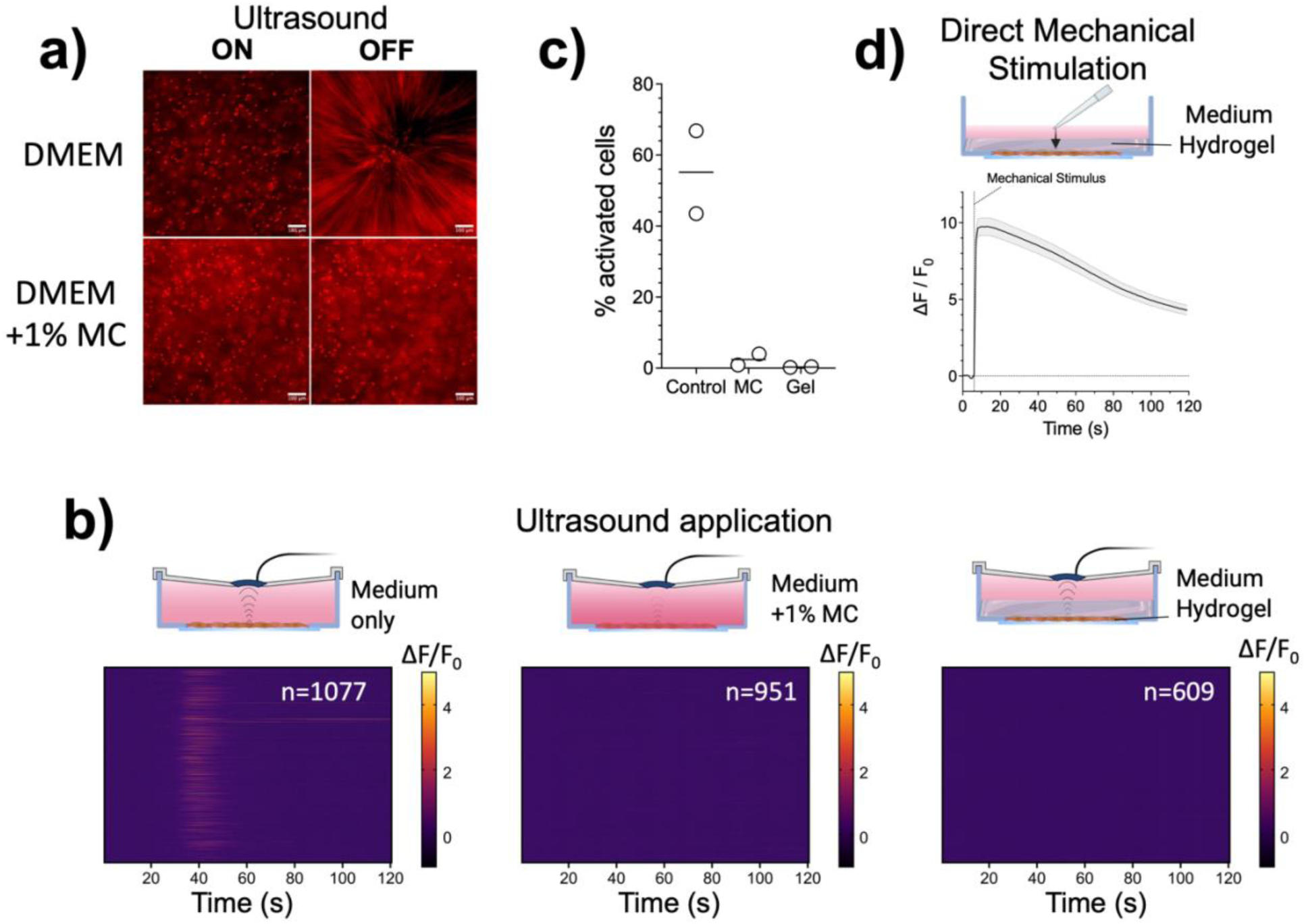
Cell response to ultrasound is inhibited when acoustic streaming is suppressed. **a)** Epi-fluorescence microscopy images (25 ms exposure time) of fluorescent particles (diameter: 5 μm) at the US focal spot before or during exposure to continuous wave ultrasound (2.2 MPa) show that inclusion of 1% methylcellulose in the supplemented culture medium greatly reduces particle movement due to acoustic streaming. **b)** Stimulation profiles of individual cells under normal conditions (supplemented DMEM), in presence of 1% methyl cellulose (MC) or after formation of a biocompatible hydrogel on top of the monolayer. **c)** The percentage of activated cells following ultrasound stimulation. Each data point corresponds to an independent experiment and the line to their average value. **E)** The average normalized fluorescence over time for NIH3T3 cells under a hydrogel, which was pressed using a pipette tip at the indicated time point. A rapid, robust Ca^2+^ was observed.

Given the above results, NIH3T3 were selected for further mechanistic experiments. The influx of calcium was not due to non-specific membrane opening, as verified by the exclusion of propidium iodide in activated cells (Figure S3). The specificity of the response was also verified by monitoring cells at a distance approximately 1 mm away from the focal spot, which did not exhibit any US-triggered Ca^2+^ influx (Figure S4). As expected, increasing sound intensity (pressure) led to higher Ca^2+^ influx (Figure 2c-e), further supporting the causal link between ultrasound application and cell activation. Accordingly, a decrease in the duty cycle or number of stimulations (pulse train repeat duration) resulted in a decrease of cell activation (Figure S5). Varying the pulse repetition frequency between 10 and 1000 Hz, did not have an appreciable effect (Figure S5).

We noted a faster response time with higher acoustic intensities (Figure 2f). Unexpectedly, however, the response times were on the order of several seconds, which is in line with some reports (Ivanovski et al., 2024; Vasan et al., 2022; Yoon et al., 2018), but contrasts others (Yoo et al., 2022). The delayed response time argues against a direct plasma membrane mechanosensitive channel opening and subsequent Ca^2+^ influx, which is expected to occur within milliseconds (Arnadottir and Chalfie, 2010; Chu et al., 2022).

It is important to note that despite reproducible trends over many independent experiments and using different NIH3T3 cell batches, variability between different cell batches and independent experiments was recorded with respect to the range and quantitative measures of activation that were obtained through our analysis pipeline (Figure S6). We considered and attempted to limit most technical sources of variation, verified regularly the sound wave intensities emitted from the employed transducers and concluded that there is an inherent biological variability in the cell response with our protocols. Therefore, in the following paragraphs, when comparing the effect of different treatments below, the results were compared to controls performed during the same measurement session and using the same batch of cells.

### 2.2 US cell stimulation is suppressed inside viscoelastic media

The slow response time between US wave application and intracellular calcium increase is not consistent with a direct mechanical activation of mechanosensitive ion channels, which are known to respond to mechanical stimuli at the microsecond to millisecond time-scale (Arnadottir and Chalfie, 2010). Instead, the delayed response pointed towards secondary effects and signaling processes that require more time. Previous studies implicated acoustic streaming, i.e. the build-up of flow driven by acoustic wave dissipation in the fluid, in cell stimulation (Liao et al., 2019; Prieto et al., 2018). In order to test the hypothesis that acoustic streaming was responsible for the observed responses, we suppressed streaming in our experimental setup by artificially increasing the viscosity of the medium by mixing in the medium 1% methyl cellulose, a biocompatible polymer often used to thicken the extracellular fluid (Buyukurganci et al., 2023). As shown in the supplementary movie S2 and Figure 2a, streaming was greatly suppressed in media containing 1% methyl cellulose. Crucially, the rise in intracellular calcium upon sonication was also dramatically decreased, with barely any cells responding (Figure 2b,c). As a complementary method, a biocompatible gelatin-based hydrogel was photo-polymerized on top of adherent cells, which were then exposed to sonication. Again, there were no signs of ultrasound-mediated cell activation, or intracellular calcium rise (Figure 2b,c). Nevertheless, the cells under the hydrogel were still responsive to mechanical stimuli, as evidenced by a rapid increase in intracellular calcium upon gentle poking of the hydrogel layer on top of the cells with a pipette tip (Figure 2d & Movie S2).

As a conclusion, our data suggest that acoustic streaming was the main physical mechanism responsible for the calcium ion influx observed on NIH3T3 cells *in vitro*.

### 2.3 Ultrasound mobilizes intracellular calcium sources through a process that requires extracellular serum components

We next focused on the origin of calcium ions that flowed into the cytoplasm and the associated mechanism downstream of ultrasonic stimulation (Figure 3a). Calcium ions could be transported from the extracellular milieu through membrane channels, some of which are mechanosensitive (Berridge et al., 2003; Jin et al., 2020). Alternatively, calcium can be released from intracellular stores, primarily from the sarcoplasmic/endoplasmic reticulum (ER) (Bagur and Hajnoczky, 2017). To block the release of calcium from the ER, we incubated cells with Thapsigargin (TG), which inhibits the sarcoplasmic/endoplasmic reticulum Ca^2+^ ATPase activity, eventually leading to depletion of intracellular Ca^2+^ stores (Lytton et al., 1991). TG-treatment essentially shut down cytoplasmic Ca^2+^ influx (Figure 3b), indicating that Ca^2+^ intracellular pools are releasing the ions in the process. Accordingly, treatment of cells with 2-APB, an inhibitor of Ca^2+^ release from the ER (Bilmen et al., 2002), completely blocked calcium transients in US-stimulated cells (Figure 3c). These data suggest that the Ca^2+^ responsible for the US-mediated rise in cytoplasmic Ca^2+^ originate from the ER.

**Figure 3.**
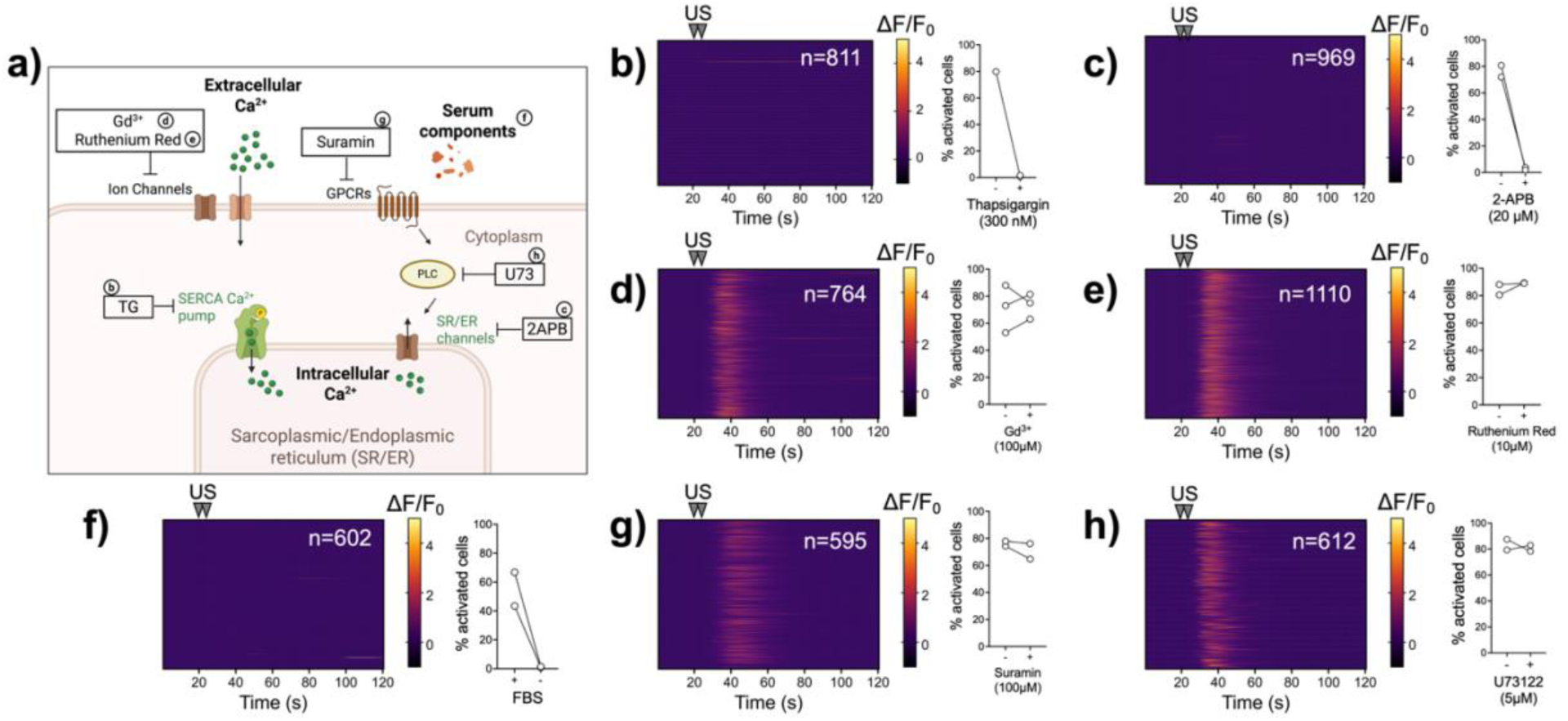
Ultrasound stimulation mobilizes intracellular calcium stores in a process that requires serum. **a)** Simplified schematic of cytoplasmic calcium influx pathways. The inhibitors used to target different components of the pathways are indicated. **b-h)** The effect of different treatments compared to control conditions. 2D histograms of normalized fluorescence increase for the treated samples from one independent experiment, and the percentage of activated cells under control and treatment conditions (2-3 independent experiments) are presented. The effect of 300 nM thapsigargin (**b**), 20 μM 2-APB (**c**), 100 μM Gd^3+^ (**d**), 10 μM ruthenium red (**e**), absence of serum (**f**), 100 μM suramin (**g**) and 5 μM U73122 (**h**) were studied. All US stimulations used the control parameters given in Table 1 and 2.2 MPa applied pressure.

The above results are also consistent with the observed latency between US application and cytoplasmic calcium increase (Figure 1f), suggesting the presence of an associated signaling pathway that triggers Ca^2+^ release from the ER, rather than direct gating of plasma membrane ion channels. Accordingly, the non-specific inhibition of mechanosensitive ion channels using Gd^3+^ ions, at a high concentration of 100 μM, did not affect the cell response (Figure 3d). A 10 μM Ruthenium Red solution, another non-specific cation channel blocker, also had no effect on cell activation following US application (Figure 3e).

In order to complement these results and elucidate the requirement of extracellular Ca^2+^, we undertook stimulation experiments leaving out sources of external Ca^2+^. First, we performed stimulations in serum-free media since serum contains Ca^2+^ ions; under serum-free conditions, calcium influx was essentially blocked (Figure 3f). This is especially surprising as the CO_2_ independent medium we used still contained 0.75 mM Ca^2+^. Even after supplementing with an additional 1 mM CaCl_2_, the cells were not responsive in the absence of serum (Movie S4). Nevertheless, the fibroblasts still had the ability to react and allow Ca^2+^ entry in serum-free medium, as evidenced by transient increases upon addition of FBS or Yoda1, an agonist of the mechanosensitive ion channel Piezo1 (Movie S5). These findings suggested that extracellular Ca^2+^ is not sufficient for the cell response to ultrasound. The results point towards either the presence of a necessary soluble factor in serum or the requirement of serum to prime cells for activation. We added FBS to previously non-responsive cells that were stimulated in serum-free medium, and monitored the cell response to US over time (Movie S6). As shown in supplementary movie S5, addition of FBS caused calcium influx when FBS was added due to the presence of stimulating factors in FBS (e.g. ATP). Therefore, the response to US was difficult to distinguish at the earliest US stimulation following FBS addition ((approximately 1 minute after addition; Figure S7). Nevertheless, the percentage of responsive cells increased rapidly and reached a plateau within 15 minutes (Figure S7). Taken together, these findings showed the necessity of yet-to-be-determined serum factors to enable US-triggered activation, and indicated that NIH3T3 fibroblasts required an adaptation phase, which lasted several minutes after serum was added, in order to become responsive.

The canonical pathway for Ca^2+^ release from the ER involves the production of inositol 1,4,5-triphosphate (IP_3_) through the action of phospholipase-C (PLC), and subsequent activation of IP_3_ receptors at the ER membrane (Berridge et al., 2000). Different PLC isoforms can be activated by G-protein coupled receptors (GPCR), many of which are mechanosensitive (Yang et al., 2013). Incubation of cells with suramin, a generic GPCR inhibitor, did not affect US-triggered calcium influx (Figure 3g). To examine the involvement of IP_3_-mediated signaling, we used the broad PLC inhibitor U73122 (Bleasdale and Fisher, 1993). Incubation of NIH3T3 fibroblasts with 5 μM U73122 did not affect Ca^2+^ influx (Figure 3h). Even at a higher U73122 concentration of 10 μM, the effect was negligible, despite a dramatic change in cell morphology (Figure S8), which is consistent with published reports on the loss of cell adhesion and cell rounding upon PLC inhibition (Jones et al., 2005) and indicated that US-induced Ca^2+^ influx does not necessitate well-adherent and spread cells.

### 2.5 US stimulation is insensitive to actin cytoskeletal disruption but sensitive to myosin II inhibition

We next turned our attention to the biophysical properties of the cells, which we hypothesized would influence their response to US by tuning the force propagation of the different mechanosensitive elements. We first tested the effects of actin cytoskeletal perturbations on NIH3T3 stimulation. Treatment with Latranculin B, which sequesters globular actin and prevents its incorporation in filaments, or Cytochalasin D, which caps actin filaments resulted in dramatic changes of the actin cytoskeleton as expected (Spector et al., 1989) (Figure 4a). In both cases, cells became rounded, and F-actin exhibited punctate staining primarily around the nuclei (Figure 4a). Surprisingly, despite the dramatic morphological changes, calcium influx was not affected by drug-induced F-actin disruption (Figure 4b,c). This finding suggested that an intact actin cytoskeleton was not necessary for ultrasound-mediated activation.

**Figure 4.**
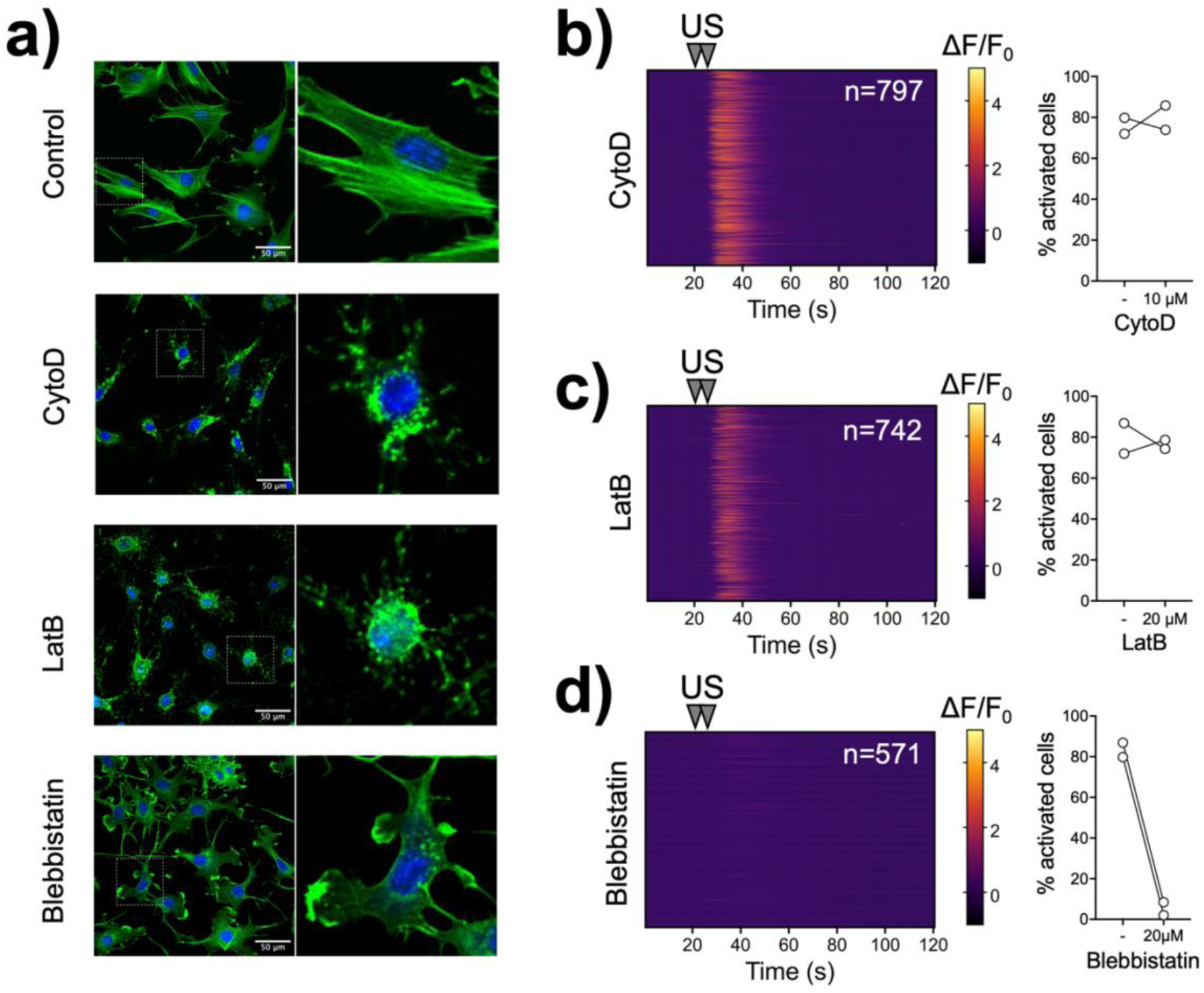
NIH3T3 fibroblast response to US is not affected by actin cytoskeleton disruption, but is abolished following myosin II inhibition. **a)** Confocal microscopy images of fixed and stained NIH3T3 fibroblasts seeded on FN-coated glass surface reveal morphological differences and actin cytoskeleton perturbations under control conditions or following 30-minute incubation with 10 μΜ cytochalasin D, 20 μΜ latranculin B or 20 μΜ blebbistatin. (F-actin: green; nuclei: blue). **b-d)** 2D histograms of normalized fluorescence increase for treated samples from one independent experiment (**b**: cytoD 10 μΜ; **c**: LatB 20 μΜ; **d**: blebbistatin 20 μΜ), as well as the percentage of activated cells under control and treatment conditions (N=2 independent experiments). US stimulations were performed using the control parameters given in Table 1 at 2.2 MPa applied pressure.

Next, the effect of the actomyosin contractility inhibitor blebbistatin was tested. Blebbistatin directly inhibits the ATPase activity of myosin II, which results in loss of contractility and a consequent reduction in cortical tension (Martens and Radmacher, 2008; Tinevez et al., 2009). Accordingly, we observed a loss of stress fibers, a flattening of cells and dendritic-like cell morphologies (Figure 4a). Intriguingly, NIH3T3 fibroblasts ceased to respond to US stimulation following blebbistatin treatment (Figure 4d). Combined, our data suggest that the effects of US on calcium influx do not require an intact actin cytoskeleton, but depend on actomyosin contractility, which we hypothesized primarily acts through modulation of cortical mechanics.

### 2.6 Changes in cortical tension affect cell-responsiveness to ultrasound stimulation

We next sought to test the hypothesis that altering the physical properties of the membrane and the cortex would affect US-stimulation calcium influx. We first depleted cholesterol using beta-cyclodextrin (bCD) to disrupt organized lipid domains and modulate membrane fluidity (Zidovetzki and Levitan, 2007). Addition of 10 mM bCD caused rapidly (within 5 minutes after addition) a drastic inhibition of calcium influx (Figure 5a). Cholesterol depletion in living cells has been reported to both increase membrane fluidity (Companyo et al., 2007; Zhang et al., 2011) and increase apparent membrane tension by enhancing the membrane-to-cortex attachment (Biswas et al., 2019; Byfield et al., 2004; Kwik et al., 2003); however, its effects are manifold and direct cholesterol-protein interactions have been reported and implicated in mechanosensitive channel regulation (Beverley and Levitan, 2024). An alternative way to fluidize the membrane and reduce membrane tension is to employ small amphiphilic molecules such as benzyl alcohol, which partition inside the lipid bilayer (Friedlander et al., 1987; Gordon et al., 1980). Incubation of NIH3T3 fibroblasts with 0.5% benzyl alcohol greatly reduced US-triggered calcium influx (Figure 5b). Importantly this effect was reversible, since washout of benzyl alcohol resulted in recovery of responsiveness to US application (Movie S7 and Figure S9. Taken together, these treatments are in line with the hypothesis that membrane biomechanical properties regulate US-triggered calcium influx.

**Figure 5.**
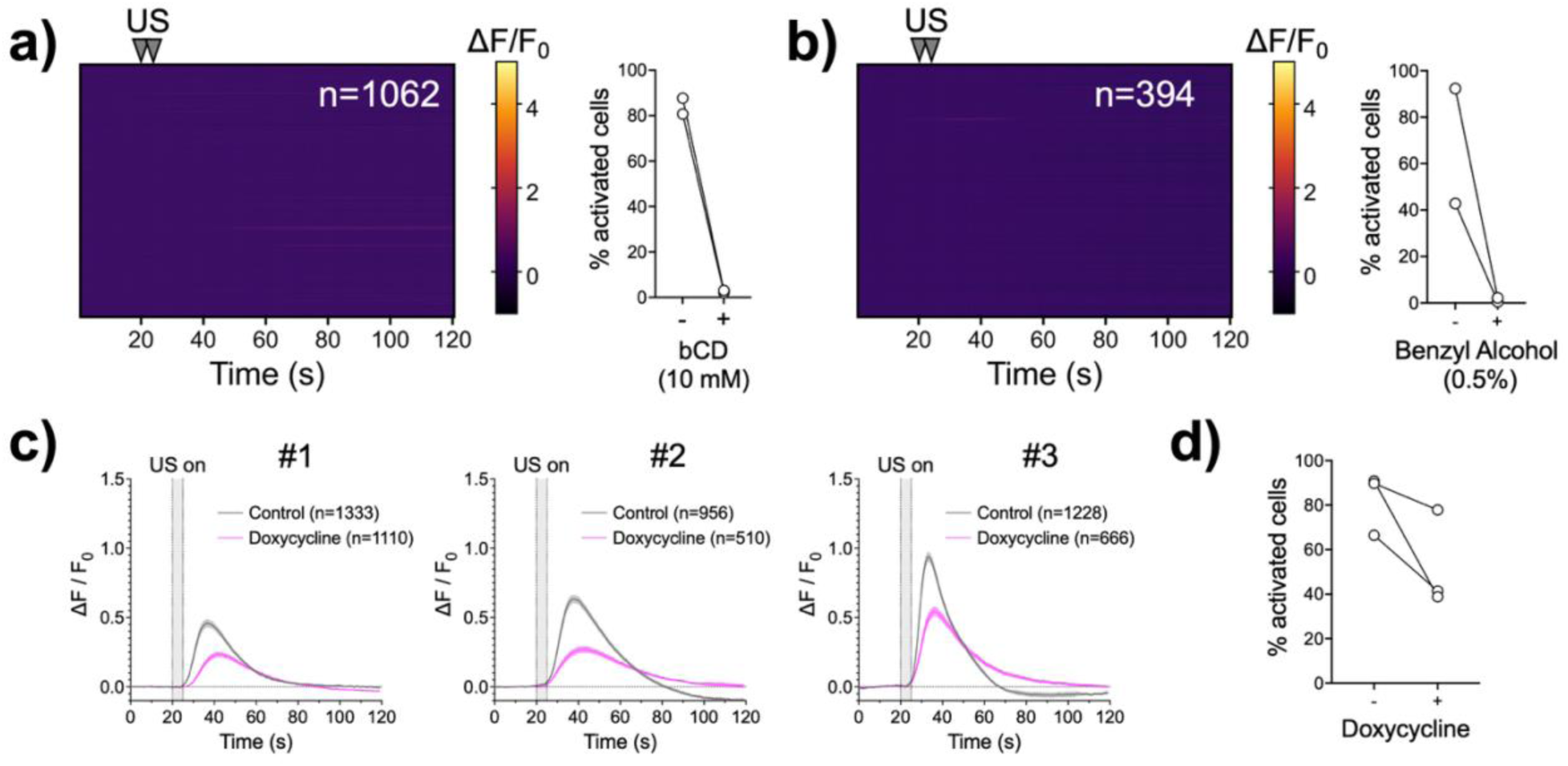
Altering the biophysical properties of the cortex through cholesterol depletion, membrane fluidization or enhancement of membrane-to-cortex attachment modulates the observed US-triggered calcium influx. **a-b)** 2D histograms of normalized fluorescence increase from one independent experiment for samples treated with 10 mM bCD (**a**) and 0.5% benzyl alcohol (**b**), and the percentage of activated cells under control and treatment conditions (N=2 independent experiments). **c)** Normalized fluorescence intensity change as a function of time (solid lines represent the average of all cells and shaded area the 95% confidence interval) for 3 independent experiments. 3T3-CAE cells were treated with 1 μg/ml doxycycline for 36 hours to induce expression of CA-Ezrin, or left untreated (control) prior to the US stimulation experiments. **d)** The percentage of activated cells under control and doxycycline-treated conditions (N=3 independent experiments). US stimulations were performed using the control parameters given in Table 1 at an applied pressure of 2.2 MPa.

In order to modulate the properties of the cortex in a more subtle manner, we tested the effect of altering the membrane-to-cortex attachment (MCA) in genetically-modified cells. The plasma membrane is coupled to the underlying actin cortex through specialized proteins, of which the ezrin-radixin-moesin (ERM) protein family is the most abundant and well-studied (Fehon et al., 2010). To increase the MCA, we induced the expression of a constitutively-active form of ezrin (CA-Ezrin), the phosphomimetic mutant T567D, using a doxycycline-inducible promoter in 3T3 fibroblasts (Bergert et al., 2021; Matsui et al., 1998). Compared to control cells, doxycycline-treated cells consistently showed decreased response in terms of the percentage of activated cells, as well as the mean fluorescence intensity rise (Movie S8 & Figure 5c,d). These data further support the hypothesis that the biophysical properties of the cortex, and in particular cortical tension, are key regulators in ultrasound-mediated cell activation.

## 3. Discussion

In this work we showed that ultrasound application on adherent cells produces a stimulus-specific, cytoplasmic calcium surge that is dependent on the cell type and the biophysical properties of its cortex. Moreover, we identified acoustic streaming as the predominant mechanism of cell activation. Surprisingly, we traced the origin of calcium ions that enter the cytoplasm to intracellular sources and ruled out the direct involvement of ion channels in the US-triggered response.

Prior research addressing the mechanisms of US-triggered cell activation *in vitro* has yielded highly heterogeneous findings, at times presenting apparent contradictions. The involvement of both mechanosensitive channels such as Piezo1 (Prieto et al., 2018; Qiu et al., 2019; Zhu et al., 2023), TRPA1 (Duque et al., 2022; Oh et al., 2019; Vasan et al., 2022), TRPC6 (Matsushita et al., 2024) and TRPP1/2 & TRPC1 (Yoo et al., 2022), as well as larger pore channels as pannexins (Lee et al., 2020; Yoon et al., 2026) and connexins (Yoon et al., 2018) has been reported. Our data indicated that mechanosensitive channels were not necessary for the observed cytoplasmic calcium influx. One plausible reason for such diverse outcomes is the use of different cell types. Indeed, we show here that the response to US was highly dependent on the cell type used, with fibroblasts showing the higher response rates. It is yet unclear how much the different biophysical cell properties vis-à-vis the different protein expression profiles are governing the variations between cell types. Nevertheless, the strong effects, that essentially switch off the response upon perturbations targeting the cortical and plasma membrane properties, argue that these properties are highly relevant and must be considered.

Another confounding factor is that the effects of pharmacological or genetic treatments to target different channels on cell biomechanics could potentially concurrently affect the mechanosensitivity or biophysical properties of cells in addition to a direct blocking of channel activity. As we have shown, changes in actomyosin contractility or membrane fluidity can have a dramatic effect on US-mediated responses, and therefore a potential shift in these parameters as a consequence of channel inhibition could indirectly lead to reduced US-triggered calcium influx. For example, Piezo1 silencing decreases the traction forces of fibroblasts (Chen et al., 2023) and alters cell mechanosensitivity (Oliva et al., 2025), potentially rendering cells non-responsive due to altered actomyosin state rather than direct ion channel transport activity.

Our findings revealed acoustic streaming as the dominant mechanism for cell activation. This second order effect of US waves leads to the build-up of fluid flow and depends on the geometry of the used setup, another potential explanation for observed differences between studies. In an elegant demonstration of how acoustic streaming affected US-triggered cell activation, Prieto et al. positioned cells at different heights from a boundary surface (Prieto et al., 2018). Acoustic streaming was dependent on the height, with no streaming at the boundary surface, while the acoustic forces were comparable: cell activation was found to be dependent on the height above the boundary, correlating with the extent of streaming. Acoustic streaming is also expected to propagate radially beyond the US focal spot; indeed, we (Figure 1c) and others (e.g Figure 1 in Yoon et al.(Yoon et al., 2026)) have recorded robust activation well beyond the focal spot, further supporting the notion that acoustic streaming constitutes a mechanism of cell activation.

The physiological relevance of US-triggered acoustic streaming *in vivo* remains understudied. A clinical study reported on use of acoustic streaming for diagnostic purposes in cysts located in breast tissue (Nightingale et al., 1999) and theoretical work (Raghavan, 2018) suggests that acoustic streaming can underlie the enhanced convective delivery of macromolecules observed in tissues (Lewis et al., 2012; Liu et al., 2010). However, the hypothesis that acoustic streaming could be used to stimulate cells *in vivo* remains untested despite indications it could play a role. For example, in the case of astrocytes, which were shown to respond to physiologically-relevant glymphatic flows that occur at the perivascular space of the brain, where the astrocyte endfeet reside (Causemann et al., 2026; Cibelli et al., 2024).

One limitation of our study is that the exact mechanism(s) of how acoustic streaming is sensed by cells and eventually leads to mobilization of intracellular calcium was not fully elucidated. Nevertheless, we provide evidence that argue against the involvement of GPCR-mediated PLC activation and subsequent IP3 generation, i.e. the canonical pathway for triggered calcium release from the ER. In addition, we highlighted the unappreciated requirement of serum for the observed response; our ongoing work tackles the challenge to identify which serum components and factors present in serum – or their combination – is necessary for sensitizing cells to US.

In summary, our results provide valuable insight on the mechanisms of ultrasound stimulation *in vitro*. In the case of fibroblasts, cells that are involved in physiological healing processes and pathological fibrotic conditions, we have identified a surprising correlation between their biophysical properties and responsiveness to US stimulation. Future work is required to check the generality of these results and their translation to the *in vivo* context.

## 4. Experimental Section

### 4.1 Materials

### 4.2 Cell Culture

HEK 293 and HeLa cells were a kind gift from the laboratory of Prof. J. Spatz (Max Planck Institute for Medical Research, Germany). NIH 3T3 fibroblasts were purchased from ATCC. Normal human dermal fibroblasts (NHDF) were purchased from PromoCell (Heidelberg, Germany). ARPE-19 cells were a kind gift from the laboratory of Dr. S. Schnichels (University of Tübingen, Germany). The NIH 3T3 transgenic cell line stably expressing the constitutively active human ezrin (T567D) in a doxycycline-inducible manner (NIH 3T3 CA-Ezrin) (Lembo et al., 2023) was a kind gift from the laboratory of Dr. A. Diz-Munoz (EMBL, Heidelberg, Germany). All cell lines were cultured as sub-confluent monolayers in Dulbecco’s Modified Eagle Medium (DMEM) supplemented with 10% FBS and 1% Penicillin-Streptomycin inside a cell culture incubator at 37 °C with 5% CO_2_.

### 4.3 Transducer Fabrication

Ultrasound transducers were assembled by fitting and gluing a lead zirconate titanate (PZT) focusing bowl into a 3D printed mount, which aligned the focus at the center of the cultured cell layer (Figure S1). The transducer was a spherical segment of ceramic type PZ54 (CTS Ferroperm), with a radius of curvature of 10mm, outer diameter of 10mm, central hole with inner diameter of 4mm and a thickness of 0.67mm. The transducer was resonant in its thickness mode at 3 MHz. Silver electrodes were screen printed on opposing sides, and a SMA-terminated coaxial cabling was soldered onto the electrodes to make electrical contacts. The ground wire was fed through the central hole to connect to the inner electrode. The transducers were driven by sinusoidal pulse trains defined on a signal generator (Tektronix AFG1062) and amplified by a 30 W power amplifier (MiniCircuits LZY-22+).

### 4.4 Transducer Characterization

Assembled transducers were submerged in water and the pressure field was mapped in 3D using a scanning needle hydrophone. The transducer was excited 15 cycle sinusoidal pulses with a fundamental frequency of 3MHz and driving amplitude 80 mV_pp_ (before amplification). Typical transverse and axial profiles of the peak pressure Figure S1. The hydrophone was then placed at the position of the peak pressure and a calibration curve was measured to assess the focal pressure as a function of driving voltage (Figure S1). Temporal peak pressures of up to 1.6 MPa were measured in the free field. This pressure corresponds to a spatial peak pulse average intensity of *I*_*SPPA*_ = 85 *W*/*cm*^2^. The focal region was ellipsoidal with a transverse focal spot size (3dB) of 0.5 mm and an axial beam length (3dB) exceeding 3mm. The peak pressure location was at 10.6 mm from the transducer face, as determined by the time of flight of the pulse to the hydrophone. Based on analytical calculations and supporting COMSOL simulations, we expect the peak pressure experienced by cells coated onto a glass cover slip to be 2x higher than that in the free field because of reflections from the glass. Therefore, in the text, we report the expected pressure acting on the cells.

### 4.5 US stimulation experiments with live-cell microscopy

Cells were seeded in 35 mm glass-bottom confocal microscopy dishes with a 20 mm glass center (VWR® Catalog # 734-2906). The bottom glass surface was coated with fibronectin following an 1-hour incubation with a 10 μg/ml fibronectin solution in PBS, followed by 3 washes with PBS. Cells were seeded at a cell density of 4×10^4^ cells/cm^2^ and incubated for 16-24 hours prior to US stimulation experiments. In the case of the NIH3T3 CA-Ezrin cells, the seeding density was 2×10^4^ cells/ml and doxycycline (1 μg/ml) was added to induce expression of the constitutively active ezrin 1 hour after cell seeding. The cells were examined 36 hours after cell seeding.

To visualize intracellular calcium levels, cells were loaded with 3μM Cal-520® AM in supplemented DMEM for 1-2 hours at 37°C. After incubation, cells were washed once and incubated with supplemented DMEM for 10 minutes to allow de-esterification of the internalized dye. Next, the medium was replaced with supplemented CO2-independent medium and the different inhibitors/chemicals were added prior to the US stimulation experiment as indicated (Table 3). Cells were imaged between 30 minutes and 4 hours from the moment of CO_2_-independent medium addition.

The 3D printed cap with the mounted ultrasound transducer was placed on top of the dishes and sealed on the sides using Teflon tape. Medium (typically 8 ml) was added to ensure immersion of the transducer in liquid. The dish was then placed on the microscope stage of a DMi8 inverted epifluorescence microscope, equipped with a heating chamber and motorized stage (Leica Microsystems). A tile scan was first acquired to determine the center of the dish, where the US focal spot is located. Time-lapse fluorescence imaging was performed for 120 seconds to monitor calcium dynamics at a frame rate of 1 frame per second (1 Hz). The US pulse train was applied at the 20 seconds time point.

### 4.6 US stimulation experiment with hydrogel on top of cells

Gelatin acryloyl was prepared as described elsewhere (Yoon et al., 2016). Following cell seeding and Cal520 AM loading, the medium was aspirated from the dish and a precursor gel solution (500 ul) containing 5.0 % w/v gelatin acryloyl and 2.6% w/v LAP was pipetted on top. The dish was then illuminated for 30 seconds using a 405 nm LED at 50 mW/cm^2^. Immediately after, supplemented CO_2_ independent medium was added on top of the gel and incubated for at least 30 minutes prior to the US stimulation experiment.

### 4.7 Image analysis

The images from the time-lapse videos were analyzed using Fiji (ImageJ) and a custom-written macro to obtain the fluorescence intensity time profiles (Figure S2). In brief, after background correction, regions of interest (ROIs) outlining individual cells were segmented and used to quantify fluorescence intensity for each frame. The resulting data were then further analyzed in Python (code available upon demand) to obtain the normalized fold-increase in dye fluorescence intensity (ΔF/F_0_). For each ROI, first the baseline value *F*_0_ before sonication (between the 3-19 second timepoints) was calculated and then the normalized ratio changes over time, 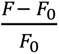, was determined. Due to stochastic intensity fluctuations (noise) we picked a threshold value with a normalized intensity ratio change greater than 30% above background to calculate the number of activated cells. The peak intensity change was also calculated for the population of activated cells. Data are presented as 2D histograms where each line corresponds to a ROI.

### 4.8 Confocal microscopy evaluation of actin cytoskeleton

NIH3T3 cells were seeded at a density of 1×10^4^ cells/cm^2^ in 8-well chambered coverglass (Nunc® Lab-Tek®) that were coated with 10 μg/ml fibronectin and culture overnight. The following day, cells were treated with inhibitors (Table 2) for 30 minutes, then washed once with PBS and fixed with 4% PFA for 15 minutes. Fixed cells were then washed 3 times with PBS and incubated with 10 μg/ml phalloidin-TRITC and 2 μg/ml DAPI in PBS for 1 hour at room temperature. Cells were washed 3 times with PBS and imaged using a laser scanning confocal microscope (Zeiss LSM880) equipped with a 20x/0.7 NA objective.

**Table 2.** List of reagents, chemicals, buffers and media used in this study.

| Chemical | Abbreviation | Company | Catalog # |
| --- | --- | --- | --- |
| Phosphate Buffer Saline | PBS | Gibco | 14190 |
| Cal-520 AM® | - | AAT Bioquest | 21130 |
| Fetal Bovine Serum | FBS | Thermo Fisher Scientific | A5209402 |
| Penicillin-Streptomycin | P/S | Thermo Fisher Scientific | 15140122 |
| Dulbecco's modified eagle medium | DMEM | Thermo Fisher Scientific | 41966 |
| CO<sub>2</sub> independent medium |  | Thermo Fisher Scientific | 18045054 |
| Methyl Cellulose | MC | Merck | M0512 |
| Gadolinium(III) chloride hexahydrate | Gd<sup>3+</sup> | Merck | G7532 |
| Suramin | - | Merck | 574625 |
| Thapsigargin | TG | Merck | 586005 |
| U73122 | - | Targetmol | T6243 |
| 2-Aminoethyl diphenylborinate | 2-APB | MedChem | HY-W009724 |
| lithium phenyl-2,4,6-trimethylbenzoylphosphinate | LAP | Merck | 900889 |
| Beta CycloDextrin | bCD | Merck | C4805 |
| Ruthenium Red | RR | Cayman Chemical | 14339 |
| Bovine Plasma Fibronectin | FN | Sigma | F1141 |
| Propidium Iodide solution | PI | Merck | P4864 |
| 5um fluorescent particle | - | Microparticle GmbH | PS-FluoRot |
| 4',6-Diamidino-2-phenylindol | DAPI | Thermo Fisher Scientific | D1306 |
| Phalloidin-tetramethylrhodamine B-isothiocyanate | Phalloidin-TRITC | Merck | P1951 |
| Blebbistatin | - | Merck | B0560 |
| Cytochalasin D | CytoD | Merck | C8273 |
| Latrunculin B | LatB | Merck | 428020 |
| Benzyl Alcohol | - | Merck | 108006 |
| Doxycycline | Dox | Merck | D9891 |
| 4% Paraformaldehyde | PFA | Thermo Fisher Scientific | J61899.AP |
| Yoda1 | - | Tocris | 5586 |

**Table 3.** List of inhibitors and treatments tested with US stimulation experiments.

| **Inhibitor/Treatment** | **Concentration** | **Incubation Time**<br>**(minutes)** | **Target** |
| --- | --- | --- | --- |
| Propidium Iodide | 10 µg/ml | >30 | Permeabilized cells |
| MC | 1% | >30 | Extracellular Viscosity |
| TG | 300 nM | 15 | Intracellular Ca<sup>2+</sup> release |
| 2-APB | 20 µM | 20-60 | Intracellular Ca<sup>2+</sup> release |
| Gd<sup>3+</sup> ions | 100 µM | 15 | Membrane cation channels |
| Ruthenium Red | 10 µM | 10 | Membrane cation channels |
| Serum-Free | - | >30 | Soluble serum factors |
| Suramin | 100 µM | 10 | GPCR inhibition |
| U73122 | 5-10 µM | 10 | Phospholipase C inhibition |
| CytoD | 10 µM | 30 | Actin cytoskeleton |
| LatB | 10 µM | 20 | Actin cytoskeleton |
| Blebbistatin | 20 µM | 20 | Actomyosin contractility |
| bCD | 10 mM | 15-30 | Cholesterol depletion |
| Benzyl Alcohol | 0.5% | 10 | Plasma Membrane Fluidity |

## 5. Acknowledgments

The authors thank Alba Diz-Munoz and Martin Bergert from the European Molecular Biology Laboratory in Heidelberg, Germany for sharing cell lines and for fruitful discussions. The authors thank Joachim Spatz from the Max Planck Institute for Medical Research in Heidelberg, Germany for granting access to the use of the microscope facilities in his laboratory.

## 6. Supporting Information

Supporting Figures S1-S9

Supporting Movies S1-S8

