## Supporting Figures S1-S9 for "Ultrasound-cell interactions mediated by cell cortex biomechanics"

for

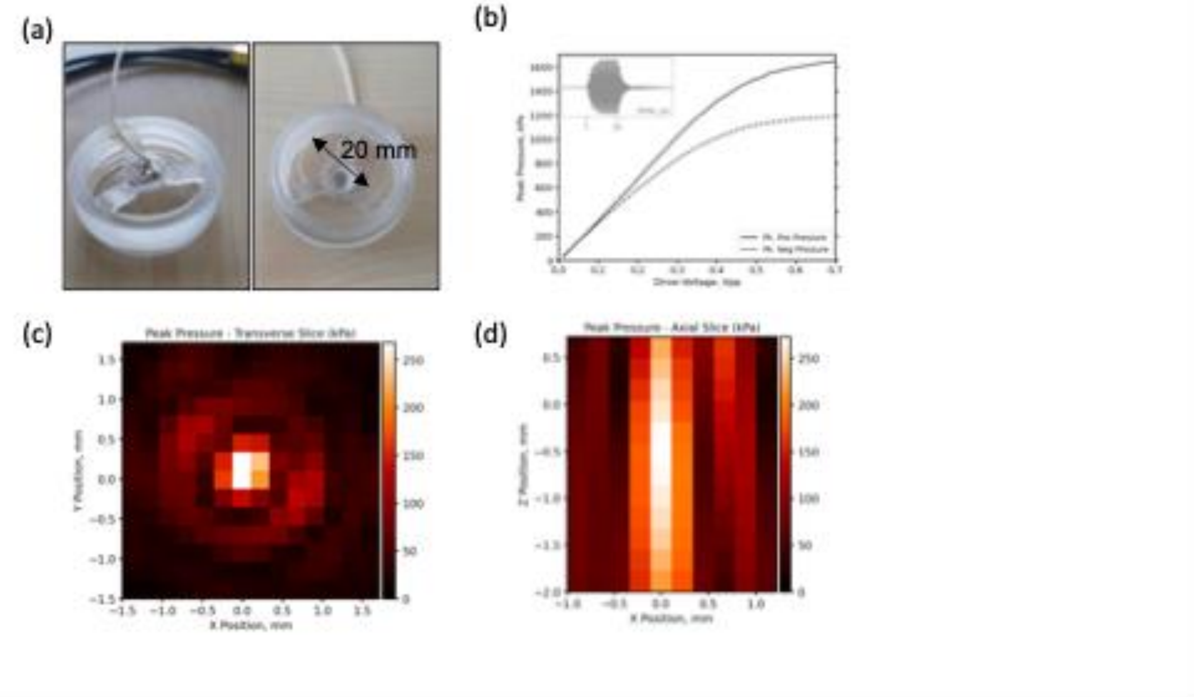


**Figure S1:** ***Transducer characterization*** **(a)** Photographs of the ultrasound transducer used in this study. The transducer was immobilized on a 3D printed mount, which could be placed on top of a commercially-available cell culture dish, that has a circular glass bottom surface of 20 mm in diameter. **(b)** Voltage calibration curve, showing the peak positive and peak negative pressure at the focus as a function of driving voltage (before amplification). Inset is a typical temporal trace of the pressure pulse, measured at 100mVpp driving. **(c)** Transverse profile of the temporal peak pressure around the focus. **(d)** Axial profile of the temporal peak pressure around the focus.

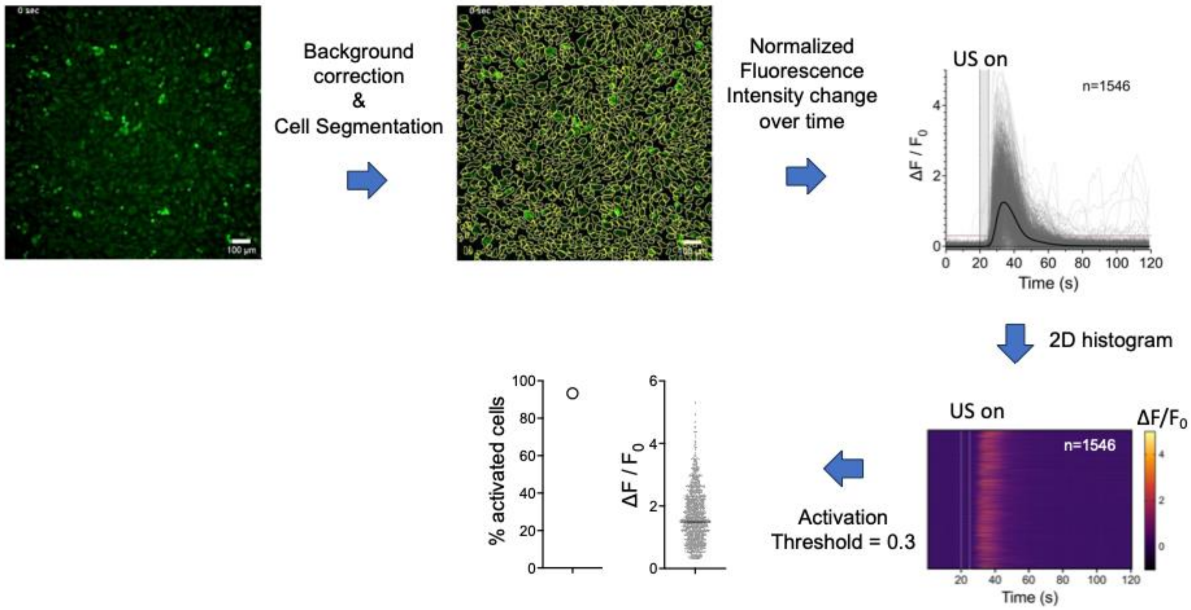


**Figure S2:** ***Image analysis pipeline description***. Time lapse videos were used to segment cells and outline ROIs that were used to calculate the normalized fluorescence intensity change over time. Data are presented as graphs of $\Delta F/F_0$ as function of time where gray lines represent individual ROIs and the solid black line the average intensity of all ROIs. Data were additionally presented as 2D histograms. By applying an activation threshold of $\Delta F/F_0$=0.3 the percentage of activated cells and the peak $\Delta F/F_0$ for the activated cells are derived.

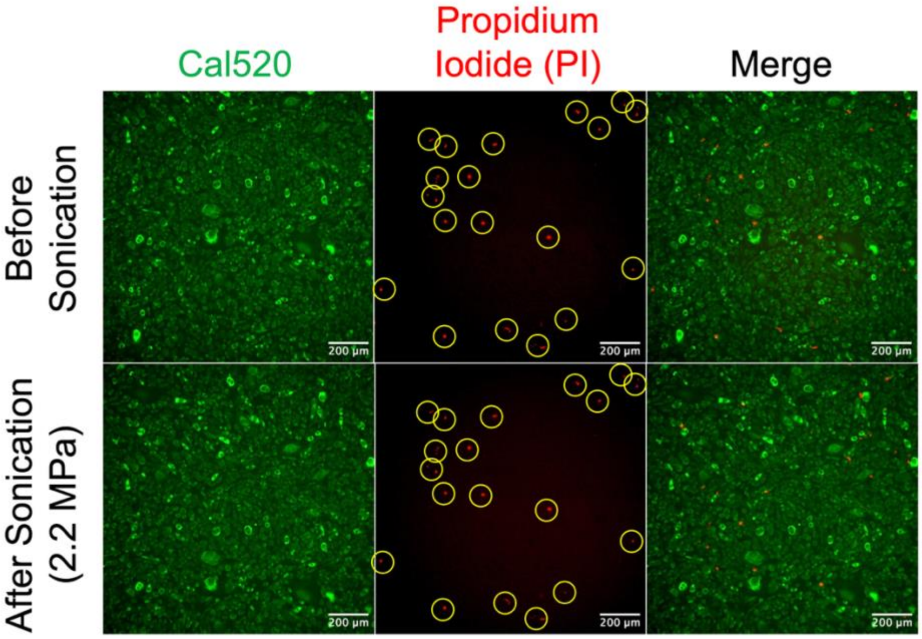


**Figure S3:** ***Propidium iodide exclusion assay***. NIH3T3 fibroblasts, loaded with Cal520AM were incubated with propidium iodide (PI; 10 µg/ml) to stain cells with permeabilized plasma membrane. PI-positive cells before sonication are already dead (membrane compromised). Application of US with the standard parameters (Table 1 in main text) and the highest intensity used in our study (2.2 MPa) did not induce non-specific openings in the plasma membrane of the cells (sonoporation). One of 3 independent experiments is shown. Yellow circles are there to denote PI-positive nuclei before sonication and are identically positioned in the image after sonication.

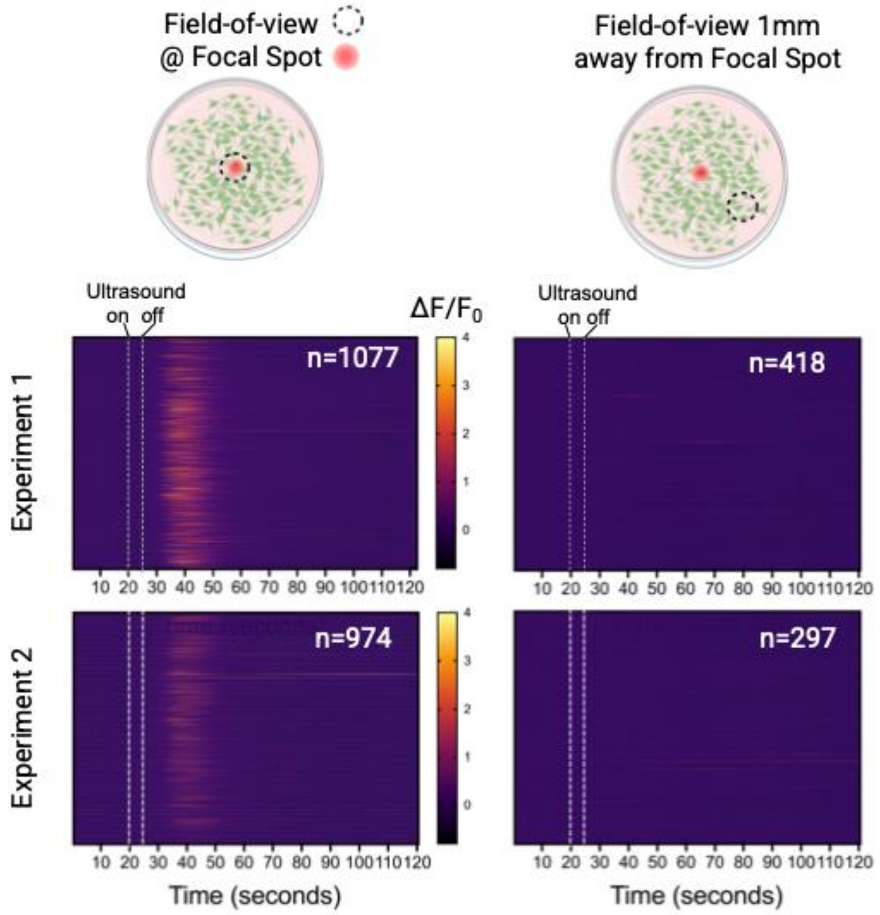


**Figure S4:** ***Ultrasound-triggered cell responses are observed specifically at the US focal spot***. 2D histograms of normalized fluorescence intensity change during US stimulation experiments recorded at the US focal spot (middle of the dish) or > 1 mm away. The transient calcium ion influx was observed in cells at the US focal spot, but not those cultured in the same dish at a distance from the US focal spot.

| | Duration | Repetition Interval/Frequency |
|---|---|---|
| Pulse | 50-20-10 ms | 100 ms / 10 Hz |
| Pulse Train | 250 ms | 500 ms/ 2 Hz |
| Pulse Train Repeat | 2.5 s | - |

| | Duration | Repetition Interval/Frequency |
|---|---|---|
| Pulse | 50 ms | 100 ms / 10 Hz |
| Pulse Train | 250 ms | 500 ms/ 2 Hz |
| Pulse Train Repeat | 2.5-0.75-0.25 s | - |

| | Duration | Repetition Interval/Frequency |
|---|---|---|
| Pulse | 50-5-0.5 ms | 100-10-1 ms / 10-100-1000 Hz |
| Pulse Train | 250 ms | 500 ms/ 2 Hz |
| Pulse Train Repeat | 2.5 s | - |

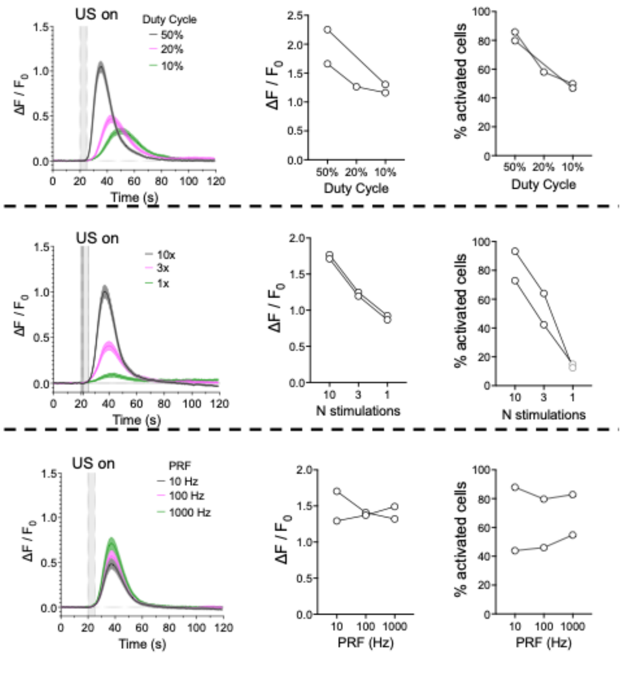


**Figure S5:** ***Dependence of US-triggered activation of NIH3T3 fibroblasts on US pulse parameters***. US stimulation of NIH3T3 fibroblasts was performed under varied parameters: duty cycle (top row), number of stimulations (middle row) and pulse repetition frequency (PRF; bottom row). The parameters according to the ITRUSST recommendations are shown at the right, with the parameter that was varied highlighted in red. The normalized change in fluorescence intensity of Cal520 (solid line represents the average of all cells and shaded area the 95% confidence interval from one of two independent experiments), the peak intensity change (each data point corresponds to the average of activated cells from an independent experiment) and the % of activated cels are presented.

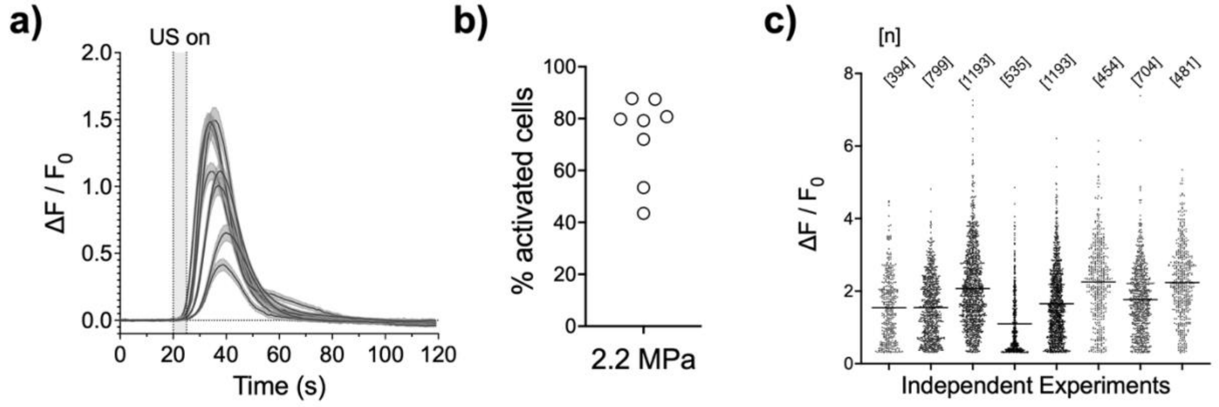


**Figure S6.** *The response of NIH3T3 fibroblasts to the standard ultrasound parameters varied between independent experiments/cell batches*. **a)** Normalized fluorescence intensity as a function of time (solid line represents the average of all cells and shaded area the 95% confidence interval), **b)** percentage of activated cells, and **c)** peak intensity change (each point corresponds to a different ROI and line represents the average value) for 8 independent experiments under the parameters shown in Table 1 of the main text and an applied pressure of 2.2 MPa.

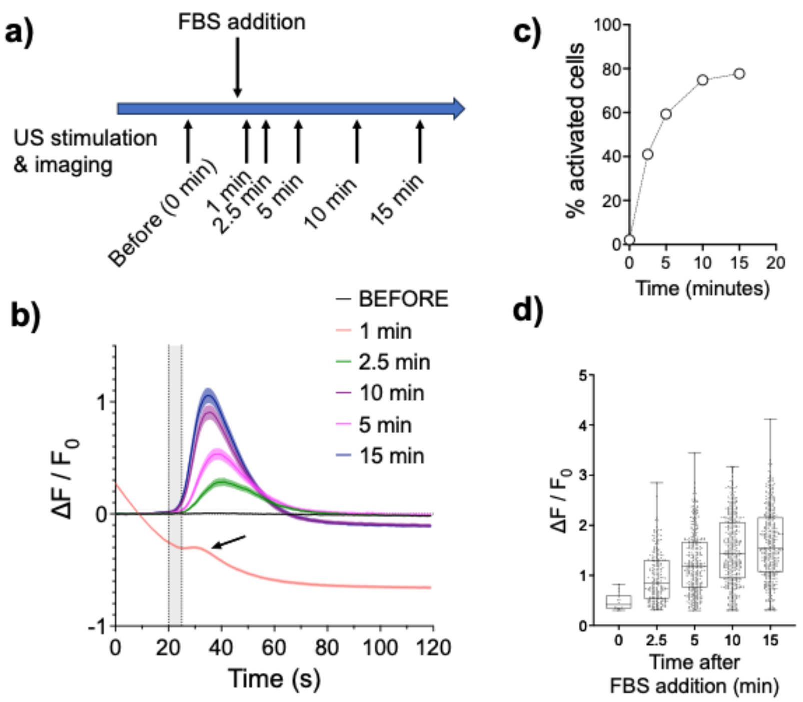


**Figure S7.** *Serum (FBS) addition to NIH3T3 fibroblasts renders the cells responsive to ultrasound in a process that takes several minutes.* **a)** Schematic of experiment to monitor US-triggered cell stimulation following FBS addition to serum-free cell culture medium. **b)** Normalized fluorescence intensity as a function of time (solid lines represent the average of all cells and shaded area the 95% confidence interval). US was applied using the control US parameters shown in Table 1 of the main text and an applied pressure of 2.2 MPa, 20 seconds after beginning of the recording (dotted lines in plot). The shape of the curve 1 minute after FBS addition is due to activation that occurs when FBS is added: cells are not yet in a background state. Nevertheless, a small peak indicates a small, but significant, US-mediated response. **c)** The percentage of activated cells at different time points following FBS addition; the time 0 corresponds to the serum-free conditions (before FBS addition). **d)** Peak fluorescence intensity at different time points following FBS addition. Each data point corresponds to a different ROI/cell, the middle line in the box the median, the box the interquartile range and whiskers the min and max values.

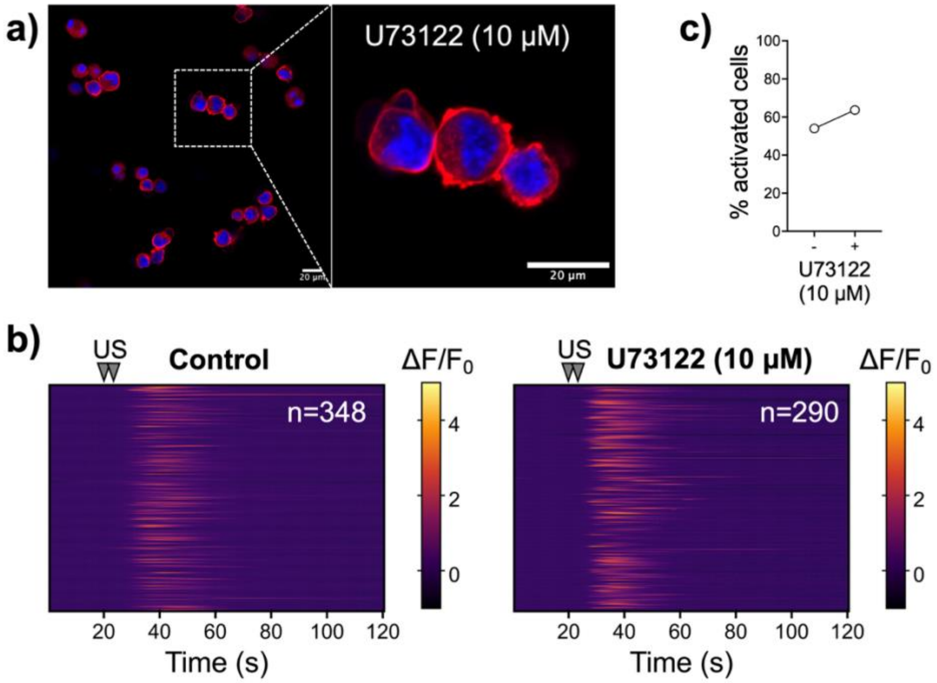


**Figure S8.** *PLC inhibition with 10 µM U73122 leads to loss of cell adhesion but does not inhibit US-triggered cell activation.* **a)** Confocal microscopy images of NIH3T3 fibroblasts seeded on FN-coated glass surface and fixed 30 minutes after incubation with 10 µM U73122. The cells were then stained against filamentous actin (TRITC-Phalloidin; red) and nuclei (DAPI; blue). Cells exhibited a spherical morphology due to loss of adhesion. **b)** 2D histograms of normalized fluorescence intensity change during US stimulation experiments recorded under control conditions or 10 minutes after treatment with 10 µM U73122. The number of cells/ROIs (n) is indicated in each case. **c)** The percentage of activated cells under control conditions, or 10 minutes after treatment with 10 µM U73122.

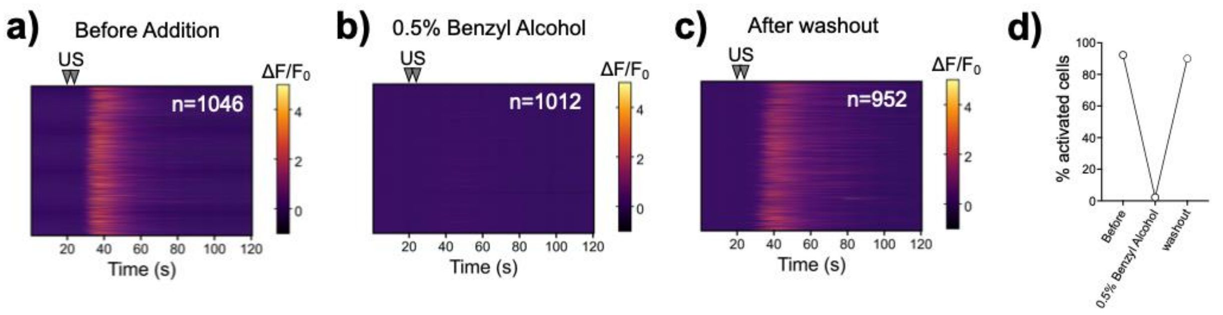


**Figure S9.** *The loss of US-triggered response of NIH3T3 fibroblasts upon incubation with benzyl alcohol is rescued after washout with fresh medium.* **a-c)** 2D histograms of normalized fluorescence intensity change during US stimulation experiments recorded before (**a**) and during incubation with 0.5% benzyl alcohol in the culture medium (**b**), as well as after exchange with fresh culture medium (**c**). **d)** The percentage of activated cells drops dramatically during a short 10-minute incubation with 0.5% benzyl alcohol, but recovers completely after the chemical is removed from the cells through exchange of medium. US stimulations were performed using the control parameters given in Table 1 of the main text at an applied pressure of 2.2 MPa.

**Supplementary Movie Legends**

**Movie S1.** *Cell type differences in response to US stimulation.* Time lapse fluorescence imaging of different Cal-520 AM-loaded cells. US stimulations were performed 20 seconds after the start of the videos, using the control parameters given in Table 1 of the main text at an applied pressure of 2.2 MPa.

**Movie S2.** *Cells are still responsive to mechanical compression when cultured under a hydrogel.* Time lapse fluorescence imaging of Cal-520 AM-loaded NIH3T3 cells under a hydrogel layer. The top of the layer was gently poked with a pipette tip to apply mechanical force 5 seconds after the start of the video.

**Movie S3.** *Addition of 1% methyl cellulose suppressed acoustic streaming.* Time lapse fluorescence imaging of fluorescent beads, 5 μm in diameter, suspended in cell culture medium with (bottom row) or without (top row) 1% methyl cellulose. An image was acquired every 36 milliseconds. A continuous US wave (3MHz; applied pressure of 2.2MPa) was applied to monitor acoustic streaming (right column). The flow observed under control conditions (top left panel; Ultrasound OFF) is due to background drift motion.

**Movie S4.** *NIH3T3 fibroblasts remain unresponsive to US in absence of serum, even in the presence of externally added calcium ions.* Time lapse fluorescence imaging of Cal-520 AM-loaded NIH3T3 cells stimulated with US in serum-free medium, supplemented with an extra 1 mM $CaCl_2$. The control US pulse train was applied 20 seconds after the start of the video with an applied pressure of 2.2 MPa.

**Movie S5.** *NIH3T3 fibroblasts cultured in serum-free media respond to addition of FBS or the Piezo1 agonist Yoda1 by calcium entry.* Time-lapse fluorescence imaging of Cal-520 AM-loaded NIH3T3 cells in serum-free medium during the addition of FBS or Yoda1.

**Movie S6.** *NIH3T3 fibroblasts gradually recover their response to US after serum is added in the culture medium.* Time lapse fluorescence imaging of Cal-520 AM-loaded NIH3T3 cells stimulated with US in serum-free medium and at different time points after the addition of FBS (the time when the time lapse was initiated is indicated). The control US pulse train was applied 20 seconds after the start of the video with an applied pressure of 2.2 MPa.

**Movie S7.** *The lack of NIH3T3 cell activation after US stimulation upon treatment with benzyl alcohol is reversible.* Time lapse fluorescence imaging of Cal-520 AM-loaded NIH3T3 cells stimulated with US before addition of benzyl alcohol (left panel), during incubation with 0.5% benzyl alcohol (middle panel) and after washout of the chemical (right panel). The control US pulse train was applied 20 seconds after the start of the video with an applied pressure of 2.2 MPa.

**Movie S8.** *Increasing the membrane-to-cortex attachment by expression of constitutively active ezrin suppressed cell activation after US stimulation.* Time lapse fluorescence imaging of Cal-520 AM-loaded NIH3T3 CA-Ezrin cells incubated with doxycycline to induce expression of constitutively active ezrin (T567D) compared with untreated cells (left panel). The control US pulse train was applied 20 seconds after the start of the video with an applied pressure of 2.2 MPa.
